# *FE DEFICIENCY-INDUCED KELCH REPEAT PROTEIN 1* (*FDK1*) links coumarin metabolism and iron homeostasis during alkaline iron deficiency in Arabidopsis

**DOI:** 10.64898/2026.10.08.757528

**Authors:** HJ Mai, C von der Mark, JC Lucas, L Schnettger, MC Knopf, EJ Stauber, M Strieker, S Metzger, V Wewer, U Wittstock, P Bauer

## Abstract

Although iron (Fe) is abundant in soils, its low solubility, particularly under alkaline conditions, often limits plant Fe acquisition. Under these conditions, coumarin-mediated Fe mobilization becomes increasingly important. *Arabidopsis thaliana FE DEFICIENCY-INDUCED KELCH REPEAT PROTEIN 1* (*FDK1*, AT3G07720) is strongly induced by Fe deficiency and regulated by the central transcription factor FIT, but its function is unknown.

We investigated *FDK1* using genetic, physiological, metabolite, and transcriptomic analyses. Loss of *FDK1* altered root growth, whereas this phenotype was restored in the *FDK1*-Ox overexpression line. Under alkaline Fe deficiency, growth medium from *FDK1-Ox* contained strongly increased levels of the Fe-mobilizing coumarins fraxetin and esculetin compared with wild type, whereas their induction was attenuated in *fdk1-1*. RNA-seq analysis linked *FDK1* abundance to coumarin biosynthesis and iron-homeostasis gene expression. *FIT* and many *FIT*-associated genes were highly expressed in *FDK1-Ox*, whereas several upstream iron-homeostasis regulators, including bHLH Ib factors, were highest in *fdk1-1*.

These findings identify *FDK1* as a component of the alkaline Fe-deficiency response that promotes iron-mobilizing coumarin accumulation and is associated with coordinated regulation of iron-homeostasis genes.

## Introduction

Iron (Fe) is an essential micronutrient for plants and a cofactor in photosynthesis, respiration, and DNA synthesis. Despite its abundance in most soils, iron is often inaccessible, particularly under alkaline conditions, where ferric iron (Fe³⁺) forms insoluble hydroxides and oxides^1,2^. Non-grass angiosperms such as *Arabidopsis thaliana* use a reduction-based iron acquisition strategy, known as Strategy I. This involves rhizosphere acidification, reduction of Fe³⁺ to Fe²⁺ by FRO2, and uptake through the high-affinity transporter IRT1, processes coordinated by the transcription factor FIT (FER-LIKE IRON DEFICIENCY INDUCED TRANSCRIPTION FACTOR)^2,3^. Under alkaline conditions, however, very low Fe³⁺ solubility limits reductive uptake, requiring additional mechanisms for iron mobilization.

One such mechanism is the biosynthesis and secretion of phenolic coumarins, including scopoletin, fraxetin, esculetin, and sideretin^4–6^. These phenylpropanoid-derived compounds are produced through enzymes including F6′H1, S8H, CYP82C4, and COSY and differ in their iron-mobilizing properties^4,6–8^. Their composition changes with environmental conditions: sideretin predominates under acidic conditions, whereas fraxetin is favored under alkaline conditions because it more effectively mobilizes iron from poorly soluble sources^5,6^. Export through the ABC transporter PDR9 is essential for efficient iron acquisition^9^, and coumarins also shape the root-associated microbiome^10,11^. Fraxetin can moreover form Fe³⁺ complexes that are taken up independently of IRT1^12^.

How coumarin biosynthesis and secretion are integrated with transcriptional iron-deficiency responses remains unclear, particularly under alkaline conditions. Specifically, the interface between FIT-dependent regulatory networks and secondary metabolism is poorly understood.

Transcriptomic analyses in *A. thaliana* have identified genes that are strongly induced by iron deficiency but remain functionally uncharacterized. Among them, *FE DEFICIENCY-INDUCED KELCH REPEAT PROTEIN 1* (*FDK1*, AT3G07720) is robustly induced under iron limitation and regulated by FIT^3,13^. Phylogenetic and transcriptomic analyses suggest homologs of this gene to be widely present across the plant kingdom^14^. In the Brassicaceae, these homologs are ancestors of a group of proteins known as specifier proteins (e.g. nitrile-specifier proteins, NSPs), i.e. enzymes involved in glucosinolate breakdown^15,16^. Iron is coordinated in the active site and represents an essential cofactor of specifier proteins^17,18^. Both FDK1 and specifier proteins are composed of Kelch domains, but so far no enzymatic activity has been demonstrated for FDK1, suggesting functional divergence^19,20^. Interestingly, specifier proteins act together with myrosinases during glucosinolate breakdown, and some of these thioglucosidases also possess β-O-glucosidase activity as demonstrated with coumarin glucosides as substrates^21^.

We investigated the role of *FDK1* in iron-deficiency responses in *A. thaliana*. We examined its subcellular localization and used knockout and overexpression lines to assess primary iron-deficiency responses and coumarin accumulation under different iron and pH conditions. Transcriptomic analyses were further employed to identify *FDK1*-associated regulatory networks and their links to coumarin metabolism and iron homeostasis.

## Results

### Loss of FDK1 alters root growth but does not strongly affect primary iron-deficiency physiology

To study the impact of FDK1 on iron deficiency responses, we used a T-DNA mutant with non-functional FDK1 identified from the SALK collection (*fdk1-1*) (Supplementary Figure 1a) and the newly generated overexpression line *FDK1-Ox* that expressed HA_3_-FDK1 under control of the 35s promoter in the *fdk1-1* background (Supplementary Figure 1b).

Seedlings were grown on solid medium for 10 days under +Fe (50 µM) or -Fe (0 µM) conditions, and root growth was assessed as a primary physiological readout. In all three genotypes, root length increased under -Fe compared to +Fe conditions (Figure 1a,b). Under both iron regimes, *fdk1-1* seedlings developed significantly longer roots than Col-0. In *FDK1-Ox*, this phenotype was complemented, with root lengths comparable to those of Col-0 under both conditions (Figure 1a,b). These observations indicate that loss of *FDK1* alters root growth, and that this phenotype is restored by *FDK1* overexpression.

**Figure 1.**
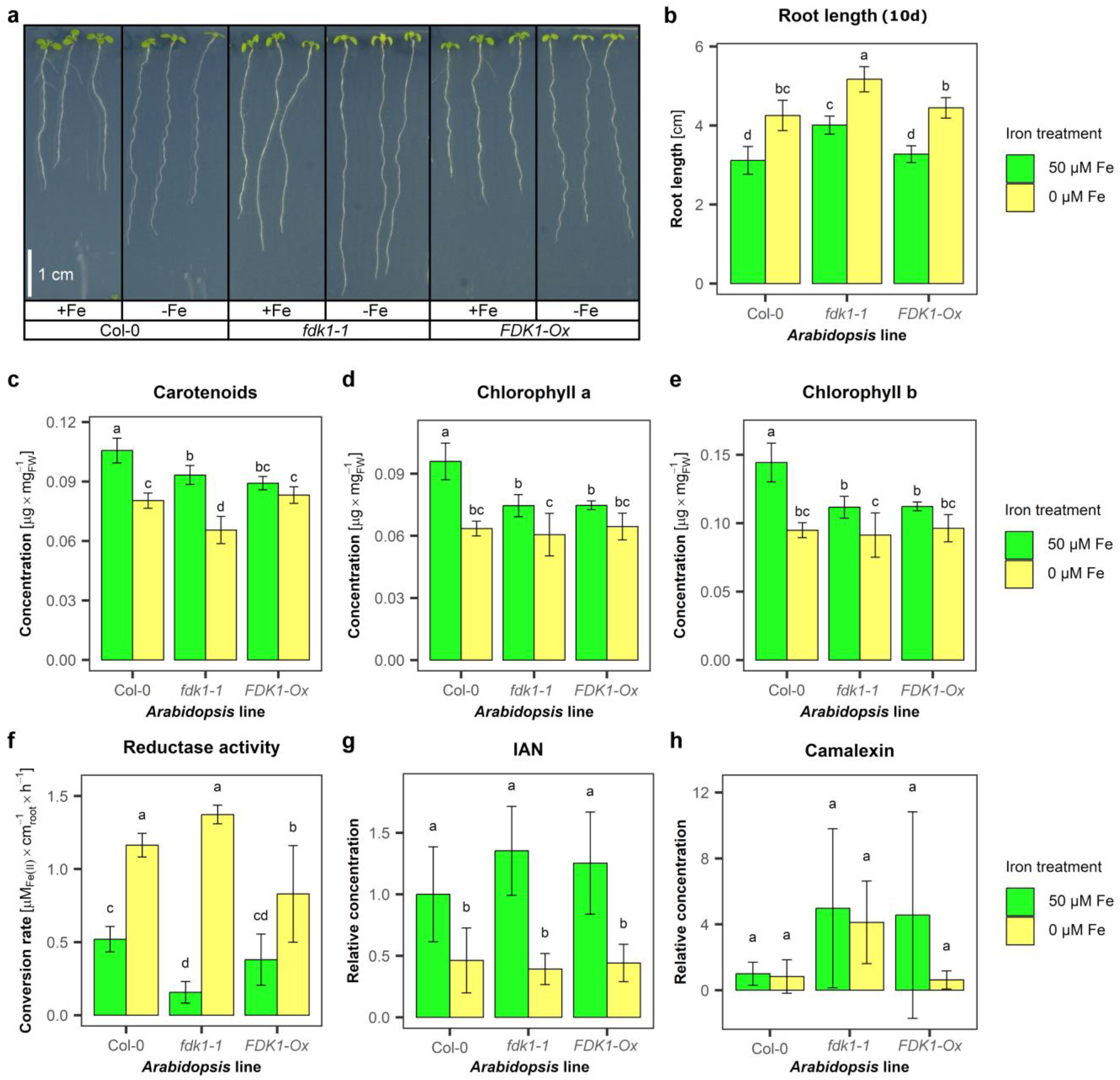
Phenotype and physiological parameters of Col-0, *fdk1-1* and *FDK1-Ox* in Arabidopsis seedlings grown on ½ Hoagland agar with (+Fe) or without iron (-Fe) for 10 days. a: root length phenotype of 10-day-old Arabidopsis seedlings, b: Root lengths, c: carotenoids contents in shoots, d: chlorophyll a content in shoots, e: chlorophyll b contents in shoots, f: reductase activity in roots, g: IAN (indole acetonitrile) relative IAN contents in roots, h: relative camalexin contents in roots. Statistical analyses were performed using ANOVA (p < 0.05) followed by Duncan’s MRT (p < 0.05). Means ± SD, N=11 (B), 3 (C, D, E, F), and 5 (G, H).

Because no clear differences in shoot coloration were observed, we next quantified carotenoid and chlorophyll contents in shoots of 10-day-old seedlings to assess whether *FDK1* affects photosynthetic pigment accumulation. Under +Fe conditions, Col-0 generally contained higher pigment levels, particularly relative to *fdk1-1* (Figure 1c-e). However, no consistent differences were detected between *fdk1-1* and *FDK1-Ox*. Under -Fe compared to +Fe, all genotypes showed reduced pigment contents, with statistically significant reductions in Col-0 and *fdk1-1* but not consistently in *FDK1-Ox* (Figure 1c-e). Overall, pigment levels did not show a clear genotype-dependent pattern that paralleled the root phenotype.

To assess whether *FDK1* influences a central component of Strategy I iron acquisition, we measured ferric reductase activity in roots of 10-day-old seedlings grown under +Fe and -Fe conditions. Reductase activity increased significantly under -Fe compared to +Fe independently of FDK1 levels (Figure 1f). Under +Fe, *fdk1-1* roots exhibited significantly lower reductase activity than Col-0, whereas under -Fe, *FDK1-Ox* roots displayed slightly lower activity than Col-0 and *fdk1-1* (Figure 1f). These differences were modest relative to the overall induction by iron deficiency.

Because FDK1 shows similarity to members of the NSP family, and indolic glucosinolate breakdown can contribute to auxin biosynthesis via indole-3-acetonitrile (IAN), we next tested whether *FDK1* affects IAN levels. In all genotypes, IAN concentrations were significantly lower under -Fe compared to +Fe (Figure 1g). However, no significant differences were detected between Col-0, *fdk1-1*, and *FDK1-Ox* under either condition. Since auxin levels were below the detection threshold in all samples, we further examined whether IAN might be redirected into camalexin biosynthesis. However, camalexin concentrations did not differ significantly between genotypes or iron treatments (Figure 1h).

Taken together, loss of *FDK1* affects root growth, whereas ferric reductase activity, pigment accumulation, IAN, and camalexin are largely unchanged across genotypes under the tested conditions.

### FDK1-GFP localizes to the cytoplasm and nucleus in *N. benthamiana* leaves and *A. thaliana* roots

To determine the subcellular localization, FDK1-GFP was transiently expressed in *Nicotiana benthamiana* leaves and stably expressed in *A. thaliana*. In tobacco leaves, FDK1-GFP localized to both the cytoplasm and nucleus (Supplementary Figure 1c). The same pattern was observed in *A. thaliana* roots under both iron-sufficient and iron-deficient conditions (Supplementary Figure 1d,e). Hence, iron availability did not visibly alter FDK1-GFP localization.

### *FDK1* is required for induction of specific coumarins under alkaline iron deficiency

Since NSPs act together with myrosinases, and some myrosinases have been shown to exhibit scopolin glucohydrolase activity^21^, we hypothesized that *FDK1* might be involved in coumarin metabolism. To test this hypothesis, we determined total phenolics and quantified individual coumarins in the growth medium. When seedlings were grown under iron-sufficient conditions at pH 5.5 or pH 7.5, total phenolic levels remained low in all genotypes (Figure 2a). In contrast, under combined iron deficiency and high pH (0 µM Fe pH 7.5), Col-0 medium showed a significant increase in phenolic content relative to both control conditions. *FDK1-Ox* also had higher phenolic levels under 0 µM Fe pH 7.5 than controls, although the increase was smaller than in Col-0. In *fdk1-1* medium, no significant increase was detected under this condition, although mean values were slightly elevated. These data show that the iron-deficiency-induced enrichment of phenolics in the medium is lacking in *fdk1-1* under alkaline iron deficiency.

**Figure 2.**
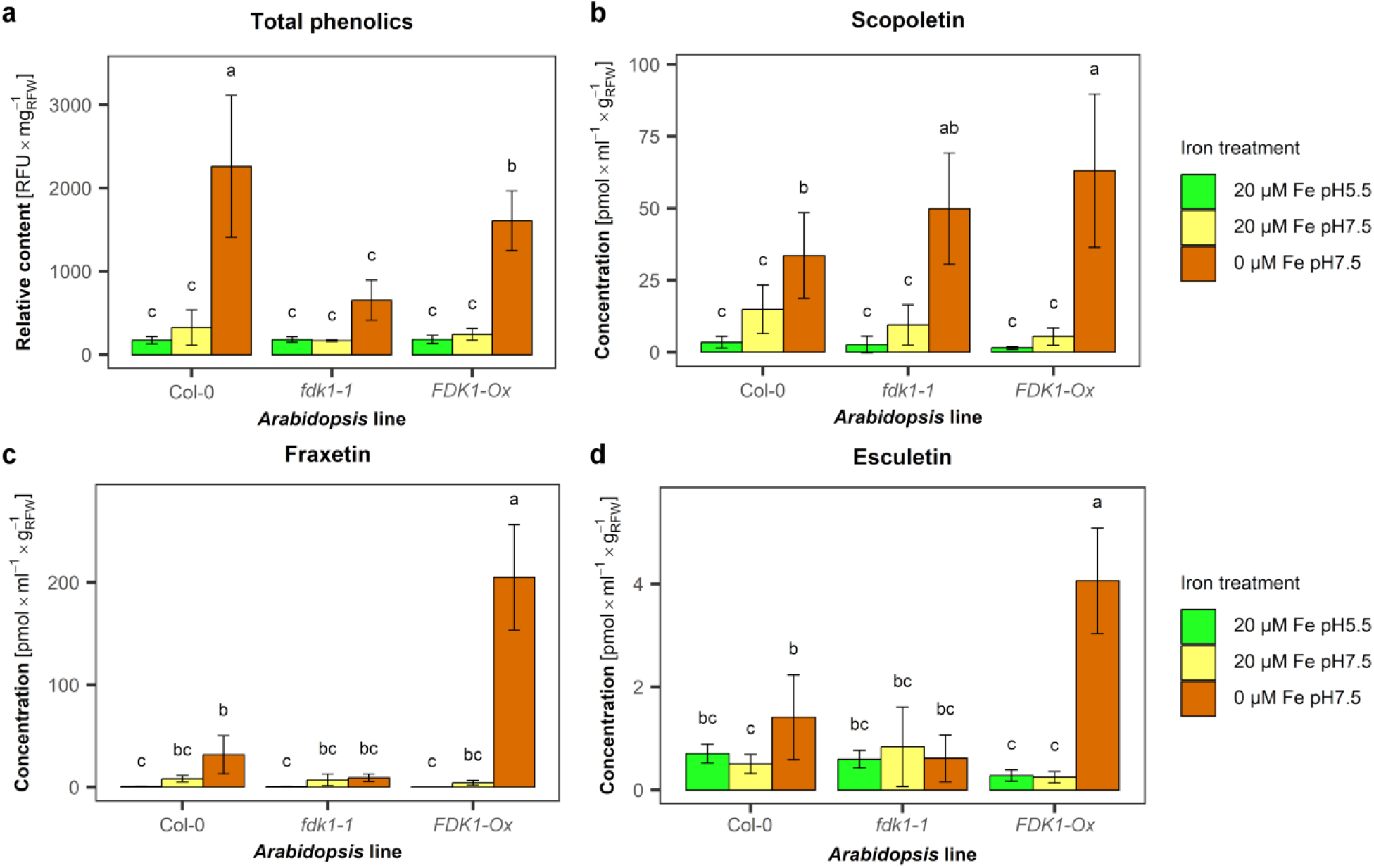
Relative total phenolics and absolute coumarin contents in the liquid medium of Arabidopsis plants grown hydroponically in liquid ½ Hoagland medium with iron at pH 5.5 (20 µM Fe pH5.5) or pH 7.5 (20 µM Fe pH7.5) or without iron at pH 7.5 (0 µM Fe pH7.5). a: Total phenolics contents in liquid medium. The seedlings were grown for 7 days on +Fe Hoagland agar plates and then transferred to a small hydroponic system with the respective iron and pH conditions for 3 days. Total phenolics were measured spectroscopically at 340 nm emission and 420 nm excitation. b-d: plants were grown hydroponically in ¼ Hoagland medium with 20 µM Fe for 5 weeks and then transferred to the respective media for 1 week. After a total of 6 weeks, the liquid medium was harvested. Fraxetin (b), scopoletin (c) and esculetin (d) contents in the medium were measured by HPLC-ESI-MS, and the results were normalized to the respective root fresh weights. Statistical analyses were performed using ANOVA (p<0.05) followed by Duncan’s MRT (p<0.05). Means ± SD, N= 3.

To obtain compound-specific information, we next quantified individual coumarins in a longer-term hydroponic experiment. Plants were grown for five weeks under iron-sufficient conditions (20 µM Fe pH 5.5) and then transferred for one week to one of three conditions: 20 µM Fe pH 5.5, 20 µM Fe pH 7.5, or 0 µM Fe pH 7.5. After this treatment, scopoletin, fraxetin, and esculetin concentrations in the growth medium were measured by HPLC-ESI-MS.

Scopoletin concentrations increased significantly under 0 µM Fe pH 7.5 in all three genotypes compared to the two iron-sufficient conditions (Figure 2b). Under this condition, *fdk1-1* did not differ significantly from Col-0, whereas *FDK1-Ox* showed higher scopoletin levels than Col-0. Thus, scopoletin levels responded to alkaline iron deficiency independently of *FDK1*, although it was enhanced in the overexpression line.

In contrast, fraxetin showed a distinct genotype-dependent pattern. In medium of Col-0, fraxetin concentrations were significantly increased under 0 µM Fe pH 7.5 compared to both iron-sufficient conditions, while no difference was observed between the two +Fe treatments (Figure 2c). In *fdk1-1* medium, fraxetin levels did not increase significantly under any condition, resulting in lower mean fraxetin concentrations compared to Col-0 under 0 µM Fe pH 7.5. In *FDK1-Ox*, fraxetin levels were strongly elevated under 0 µM Fe pH 7.5, with increases exceeding 1600-fold and 48-fold relative to 20 µM Fe pH 5.5 and 20 µM Fe pH 7.5, respectively. Under this condition, fraxetin levels in *FDK1-Ox* exceeded those in Col-0 by more than sixfold.

Esculetin displayed a similar but less pronounced pattern than fraxetin. In Col-0 medium, esculetin levels were significantly increased under 0 µM Fe pH 7.5 compared to 20 µM Fe pH 7.5, but not compared to 20 µM Fe pH 5.5 (Figure 2d). In *fdk1-1* medium, esculetin levels did not change across conditions. In *FDK1-Ox* medium, esculetin concentrations increased strongly under 0 µM Fe pH 7.5, exceeding those of the other two conditions by more than 14-fold and those of Col-0 under the same condition by more than 2.5-fold. Overall esculetin concentrations were lower than those of fraxetin and scopoletin across all treatments.

Taken together, combined alkaline pH and iron deficiency triggered the strongest coumarin response in Col-0. Under these conditions, scopoletin increased in medium of all genotypes, whereas fraxetin was elevated onlyin medium of *FDK1-Ox* and Col-0 and esculetin only in *FDK1-Ox* medium. These observations indicate that *FDK1* is required for an appropriate fraxetin and esculetin induction under alkaline iron deficiency.

### Mineral nutrient profiles are primarily determined by treatment rather than genotype

Given the pronounced differences in coumarin levels, we next asked whether these changes are reflected in plant mineral nutrient status. Across all genotypes, iron concentrations were highest under +Fe pH 5.5, reduced under -Fe pH 5.5, and lowest under -Fe pH 7.5 (Figure 3a). Differences between genotypes within a given treatment were limited. Similar treatment-dependent patterns were observed for other elements. Zinc increased under -Fe pH 5.5 in Col-0 and *fdk1-1* and was lower under -Fe pH 7.5 in all three genotypes (Figure 3b), while copper increased under iron deficiency, with *FDK1-Ox* showing higher levels than Col-0 under -Fe pH 7.5 (Figure 3c). Manganese concentrations were generally higher under -Fe pH 7.5 than under the other treatments (Figure 3d). Phenospex-based analysis detected no genotype-dependent differences among Col-0, *fdk1-1*, and *FDK1-Ox* on control or alkaline calcareous soil (Supplementary Figure 3).

**Figure 3.**
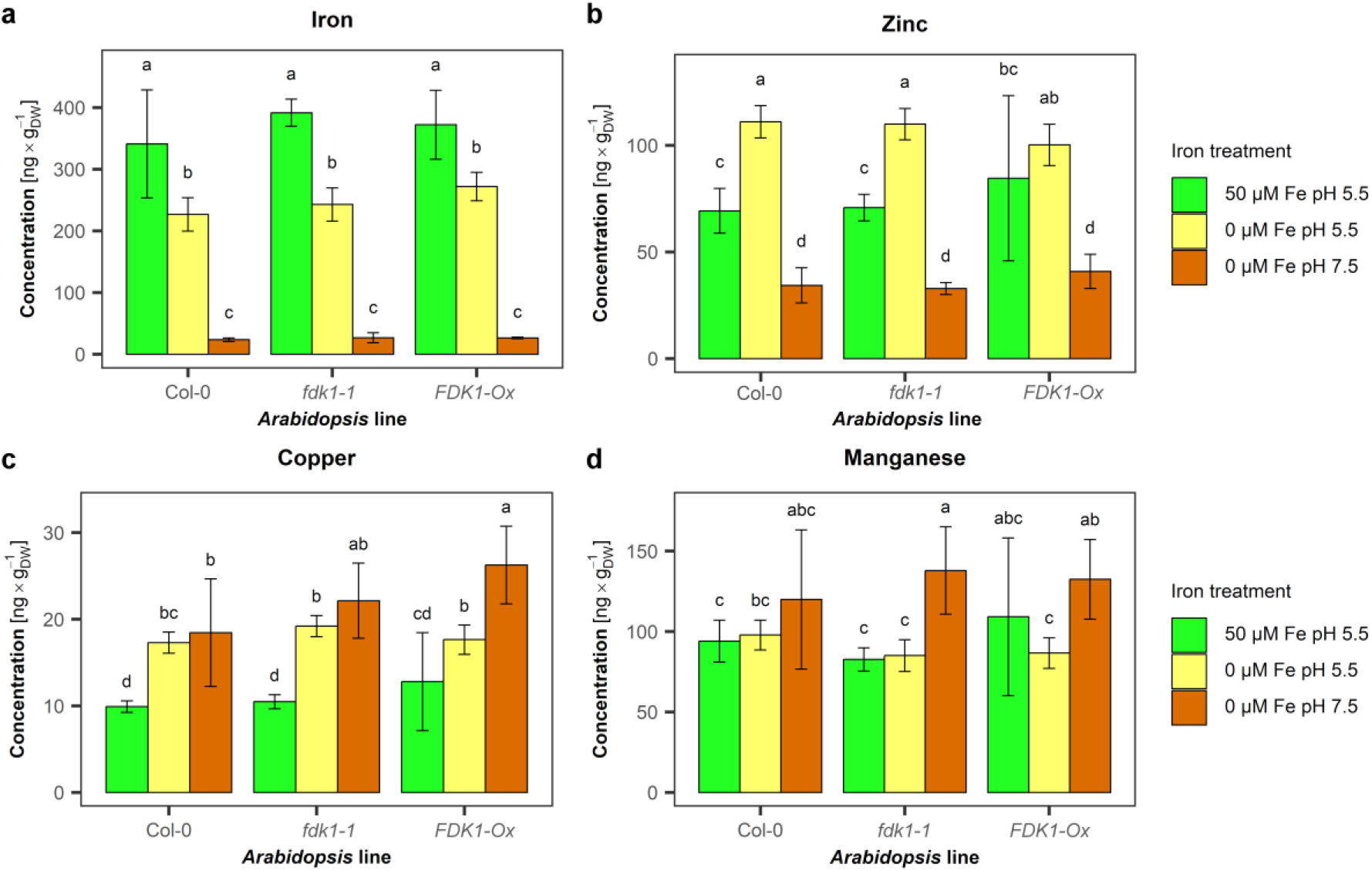
Mineral contents in shoots of 7-day-old Arabidopsis seedlings grown on ½ Hoagland agar plates with 50 µM Fe pH 5.5, 0 µM Fe pH 5.5 and 0 µM Fe pH 7.5. a: iron, b: zinc, c: copper, d: manganese. Mineral contents were measured by ICP-MS and, normalized to the dry weight of material. Statistical analyses were conducted using ANOVA (α < 0.05, p < 0.05) followed by Duncan’s MRT (α < 0.05, p < 0.05). Means ± SD, N=5.

Overall, iron deficiency and pH strongly influenced mineral accumulation, whereas genotype-dependent effects were comparatively small.

### Differentially expressed genes are enriched for GO terms related to coumarin metabolism, iron starvation, and pH-related processes

To gain a global view of the processes underlying the *FDK1*-related phenotypes and learn more about the role of *FDK1*, we next analyzed transcriptional responses across genotypes and treatments. Principal component analysis separated samples primarily by treatment along PC1, with -Fe pH 7.5 clearly distinct from the other conditions, and by genotype along PC2, where *fdk1-1* was separated from Col-0 and *FDK1-Ox* (Supplementary Figure 2).

To dissect the biological processes underlying these transcriptional changes, we analyzed the functional composition of differentially expressed genes. Gene expression analysis was conducted using a total of 18 pairwise comparisons (Supplementary Table 1). In total, 6690 genes were significantly regulated in at least one comparison. Hierarchical clustering of these genes separated them into two superclusters and 15 smaller clusters (Figure 4a, Supplementary Table 2). Supercluster 1 genes were generally expressed at lower levels under high pH conditions compared to low pH, whereas supercluster 2 genes were expressed at higher levels under high pH compared to low pH (Figure 4a, Supplementary Table 2).

**Figure 4.**
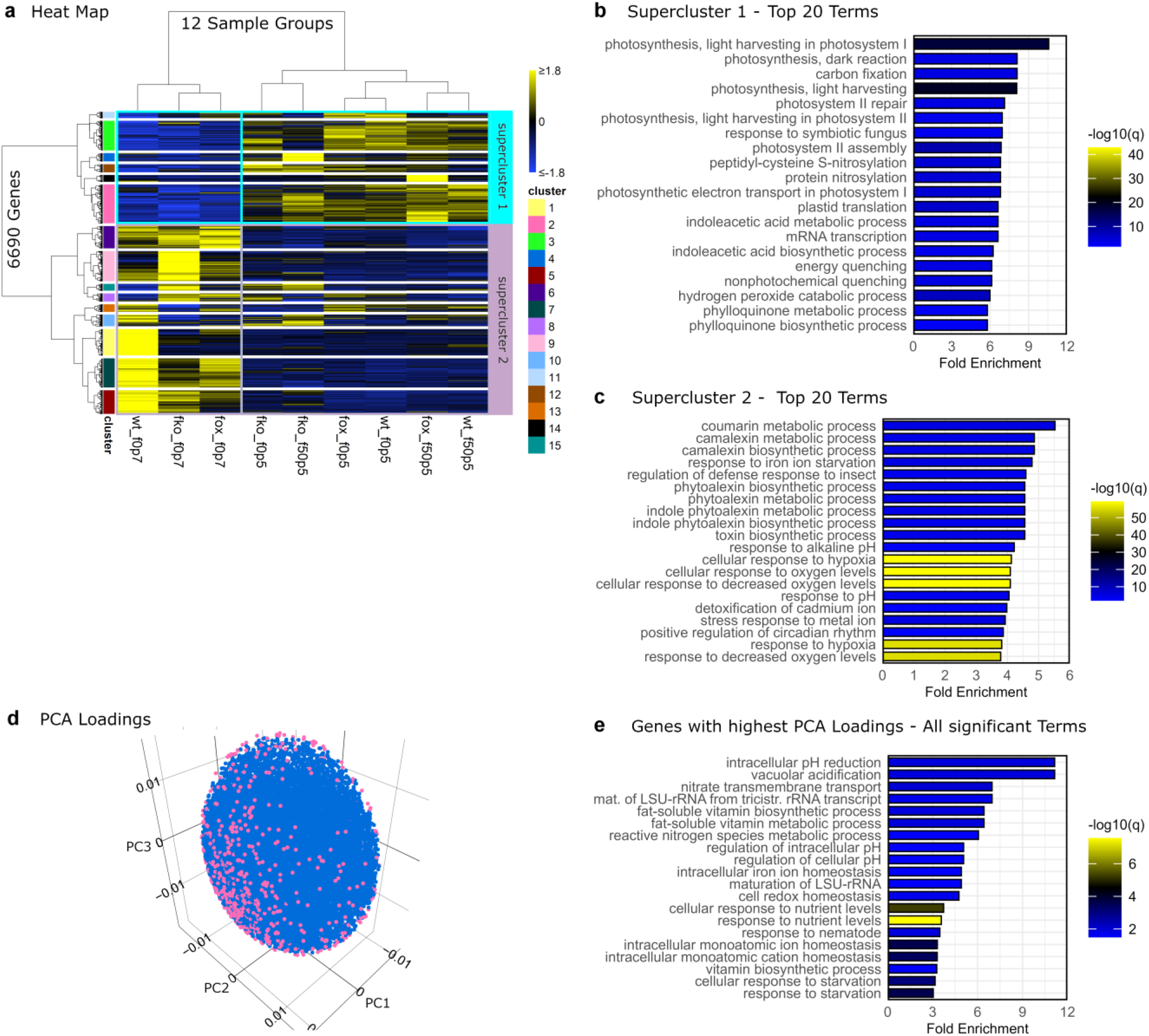
Heatmap of regulated genes, Gene Ontology (GO) term enrichment analysis of the two superclusters and GO term enrichment of genes with the strongest PCA loadings. a: Heatmap of our 12 sample groups and with genes that were regulated at least once in the 18 pairwise comparisons performed in our RNA-Seq experiment. The clustering method was Ward D2. The abbreviations of the sample groups are: wt = wildtype, fko = *fdk1-1* knockout mutant, fox = *FDK1-Ox* overexpression mutant, f50p5 = 50 µM Fe pH 5.5, f0p5 = 0 µM Fe pH 5.5, f0p7 = 0 µM Fe pH 7.5. b: GO term enrichment analysis of the supercluster 1. c: GO term enrichment analysis of the supercluster 2. d: visual representation of selected genes according to their PCA loadings. The genes were plotted in 3-D space according to their PCA loadings with PC1, PC2 and PC3 loadings as the coordinate axes. According to the spread of the loadings, an ellipsoid was constructed tightly around the data. Then the axes were iteratively shrunk until 999 genes were plotted outside the ellipsoid (pink dots). These were considered important for separation of the genotypes and treatments in PCA analysis. e: GO term enrichment analysis of the 999 genes with the highest PCA loadings. GO enrichment p-values were corrected using the Benjamini-Hochberg method.

To determine the functional characteristics of these gene sets, which may reflect the direct and indirect effects of *FDK1*, we performed GO term enrichment analysis. Supercluster 1 was dominated by genes related to photosynthesis (Figure 4b, Supplementary Table 3, Supplementary Table 4). In contrast, supercluster 2 was enriched for terms such as “coumarin metabolic process,” “camalexin metabolic process,” and “camalexin biosynthetic process,” which showed the highest fold enrichments, as well as terms related to “response to iron ion starvation” and pH-related processes (Figure 4c, Supplementary Table 5, Supplementary Table 6). These results indicate that genes induced under high pH and iron deficiency are associated with both specialized metabolism and iron starvation responses.

Because the number of regulated genes was large, we next applied a selection strategy to focus on those contributing most strongly to variation in the dataset. 999 genes were selected based on their PCA loadings, representing their contribution to principal components (Figure 4d, Supplementary Table 7). Genes with the most extreme loadings were considered as the most informative for distinguishing treatments and genotypes. GO term enrichment analysis of this subset revealed enrichment for pH-related processes, fat-soluble vitamin-related terms, and ion and iron homeostasis, as well as starvation-related processes (Figure 4e, Supplementary Table 8). These enriched terms are consistent with the imposed iron and pH treatments, indicating that the selected genes capture key biological responses under these conditions.

Together, these analyses show that iron deficiency and high pH induce coordinated transcriptional changes affecting coumarin metabolism, iron homeostasis, and stress-related processes.

### Cluster-specific expression patterns link *FDK1* transcript abundance to coumarin biosynthesis and metal homeostasis

To identify genotype-specific transcriptional signatures, we next examined individual gene clusters derived from the hierarchical clustering. Cluster 6 contained genes that were expressed at the highest levels in *FDK1-Ox* under -Fe pH 7.5 conditions (Figure 4a, Supplementary Table 2). GO term enrichment analysis showed that “coumarin metabolic process” was the term with the highest fold enrichment in this cluster (Figure 5a, Supplementary Table 9). In addition, terms such as “response to iron ion starvation,” “zinc transport,” “iron transport,” and “regulation of metal ion transport” were enriched. Notably, “transition metal transmembrane transport” and “iron ion transmembrane transport” were enriched together with “intracellular iron ion homeostasis” (Figure 5a, Supplementary Table 9). These results indicate that genes associated with coumarin metabolism and metal homeostasis are highly expressed in *FDK1-Ox* under alkaline iron deficiency.

**Figure 5.**
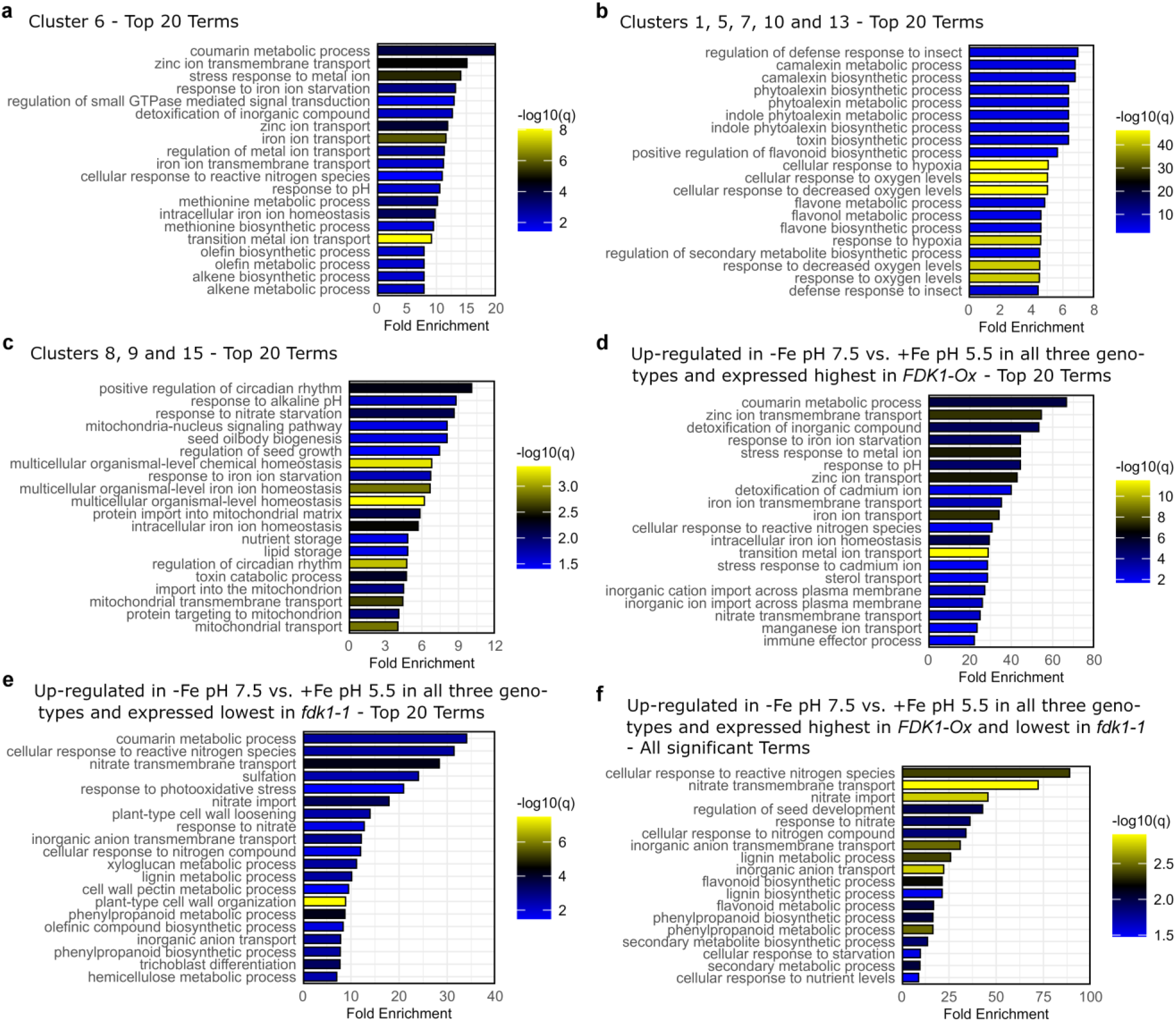
Gene Ontology (GO) term enrichment analysis of heat-map clusters derived from the RNA-Seq data, together with three manually defined gene sets. The manual sets were constructed based on expression strengths across the different genotypes under 0 µM Fe pH 7.5, with the additional requirement that all included genes were induced in all three genotypes (Col-0, *fdk1-1* and *FDK1-Ox*) under 0 µM Fe pH 7.5 compared to 50 µM Fe pH 5.5. These analyses are meant to highlight functional differences between *fdk1-1* and *FDK1-Ox* and help identify potential roles of *FDK1*. The panels show the top 20 terms except where the number of significant terms was <20. GO enrichment p-values were corrected using the Benjamini-Hochberg method.

Clusters 1, 5, 7, 10, and 13 contained genes that were most highly expressed in Col-0 under -Fe high-pH conditions (Figure 4a, Supplementary Table 2). Among the top enriched terms were indole phytoalexin- and camalexin-related processes, as well as responses to insects. In addition, oxygen-, flavone-, and flavonol-related terms were enriched (Figure 5b, Supplementary Table 10). These clusters therefore represent gene sets that are preferentially expressed in the wild type under these conditions.

Clusters 8, 9, and 15 predominantly contained genes that were most highly expressed in the *fdk1-1* knockout mutant (Figure 4a, Supplementary Table 2). These clusters were enriched for terms related to iron homeostasis and iron starvation, as well as mitochondrial transmembrane transport and protein import into mitochondria (Figure 5c, Supplementary Table 11). Thus, genes associated with iron homeostasis regulation and mitochondrial functions were preferentially expressed in *fdk1-1* under -Fe high-pH conditions.

These cluster-specific patterns distinguish the genotypes and link *FDK1* transcript abundance to coumarin metabolism, metal homeostasis, and mitochondrial processes.

### Manual gene set filtering identifies candidate *FDK1*-responsive genes under alkaline iron deficiency

Gene lists were constructed based on two criteria. First, genes had to be induced in all three genotypes under -Fe pH 7.5 compared to +Fe pH 5.5, selecting for genes generally responsive to this condition. Second, genes had to either be expressed at the highest levels in *FDK1-Ox* (Supplementary Table 12) or at the lowest levels in *fdk1-1* (Supplementary Table 13). A third list comprised genes fulfilling both criteria (Supplementary Table 14).

If these gene sets are associated with the coumarin phenotype, enrichment of coumarin-related processes would be expected. GO term enrichment analysis showed that in genes expressed highest in *FDK1-Ox*, “coumarin metabolic process” was again the term with the highest fold enrichment (Figure 5d, Supplementary Table 15). In addition, this set was enriched for zinc-, iron-, manganese-, and cadmium-related terms, many of which included transport-related processes such as “ion transport,” “transmembrane transport,” and “import across plasma membrane.”

In the gene set expressed at the lowest levels in *fdk1-1*, “coumarin metabolic process” was also the top enriched term, together with “phenylpropanoid metabolic process” and “phenylpropanoid biosynthetic process” (Figure 5e, Supplementary Table 16).

The overlapping gene set, comprising genes lowest in *fdk1-1* and highest in *FDK1-Ox*, was enriched for “cellular response to reactive nitrogen species” and “nitrate membrane transport,” as well as “phenylpropanoid metabolic process,” “phenylpropanoid biosynthetic process,” and broader secondary metabolic processes (Figure 5f, Supplementary Table 17).

### Coumarin biosynthesis genes show contrasting expression in *fdk1-1* and *FDK1-Ox*

When the three genotypes were compared under alkaline iron deficiency, *fdk1-1* had the lowest expression of several genes, including *PAL4*, *4CL3*, *CCoAOMT1*, *CCoAOMT7*, *COMT1*, and *COSY* (Table 1, Supplementary Table 1). Conversely, *PAL2*, *4CL2*, *F6’H1*, *S8H*, *BGLU42*, and *PDR9* were expressed at the highest levels in *FDK1-Ox* under the same condition. In addition, *PAL1*, *4CL1*, and *C3H* were both lowest in *fdk1-1* and highest in *FDK1-Ox* (Table 1, Supplementary Table 1).

**Table 1.**
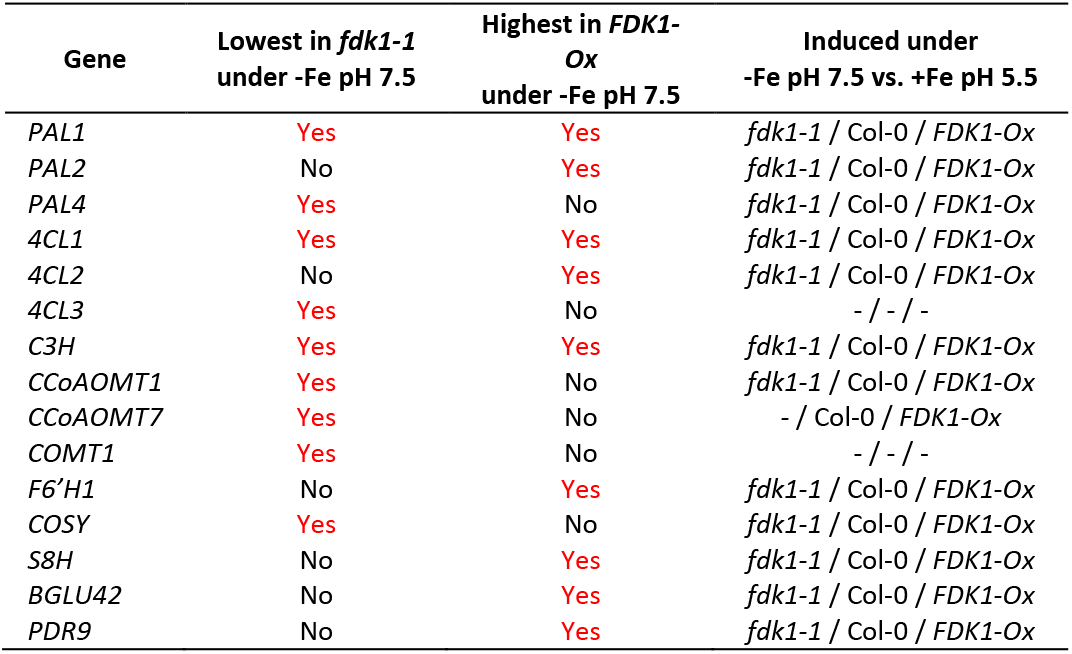
Potential coumarin biosynthesis genes that were found regulated in at least one pairwise comparison in our RNA-Seq analysis. Expression of the coumarin biosynthesis genes in the contrasting lines *fdk1-1* and *FDK1-Ox* particularly under -Fe pH 7.5 with the question whether they were expressed lowest in *fdk1-1* and/or highest in *FDK1-Ox*.

| Gene | Lowest in *fdk1-1*<br>under -Fe pH 7.5 | Highest in *FDK1-Ox*<br>under -Fe pH 7.5 | Induced under<br>-Fe pH 7.5 vs. +Fe pH 5.5 |
| --- | --- | --- | --- |
| *PAL1* | Yes | Yes | *fdk1-1* / Col-0 / *FDK1-Ox* |
| *PAL2* | No | Yes | *fdk1-1* / Col-0 / *FDK1-Ox* |
| *PAL4* | Yes | No | *fdk1-1* / Col-0 / *FDK1-Ox* |
| *4CL1* | Yes | Yes | *fdk1-1* / Col-0 / *FDK1-Ox* |
| *4CL2* | No | Yes | *fdk1-1* / Col-0 / *FDK1-Ox* |
| *4CL3* | Yes | No | - / - / - |
| *C3H* | Yes | Yes | *fdk1-1* / Col-0 / *FDK1-Ox* |
| *CCoAOMT1* | Yes | No | *fdk1-1* / Col-0 / *FDK1-Ox* |
| *CCoAOMT7* | Yes | No | - / Col-0 / *FDK1-Ox* |
| *COMT1* | Yes | No | - / - / - |
| *F6'H1* | No | Yes | *fdk1-1* / Col-0 / *FDK1-Ox* |
| *COSY* | Yes | No | *fdk1-1* / Col-0 / *FDK1-Ox* |
| *S8H* | No | Yes | *fdk1-1* / Col-0 / *FDK1-Ox* |
| *BGLU42* | No | Yes | *fdk1-1* / Col-0 / *FDK1-Ox* |
| *PDR9* | No | Yes | *fdk1-1* / Col-0 / *FDK1-Ox* |

These patterns show that multiple genes involved in coumarin biosynthesis and transport differ between genotypes under -Fe pH 7.5. Notably, *BGLU42* and *PDR9* were highly expressed in *FDK1-Ox* but not consistently lowest in *fdk1-1*, indicating that not all components follow the same pattern. Overall, expression of several coumarin biosynthesis genes differed between *fdk1-1* and *FDK1-Ox* in a manner consistent with the metabolite profiles in the culture medium.

### Iron homeostasis genes reveal contrasting FIT- and bHLH-associated expression patterns

To place these findings in the broader regulatory context of iron homeostasis, we analyzed genes from the “extended 2” iron homeostasis gene list^22^. Of the 161 genes in this list, 93 were differentially expressed in at least one comparison. These genes were distributed across several clusters (Figure 6, Supplementary Table 18). One cluster contained ferritin genes (*FER1*-*FER4*) and VIT-LIKE transporters (*VTL1*, *VTL2*, *VTL5*). Another cluster, which was the largest, contained *FIT* and many genes previously associated with FIT-dependent responses^23^, including *IRT1*, *FRO2*, *COPT2*, *F6’H1*, *S8H*, *HMA3*, *MTP8*, *PDR9*, *BGLU42*, *BTSL2*, *MYB10*, and *MYB72*. These genes were expressed at the highest levels in *FDK1-Ox* under -Fe pH 7.5, followed by *fdk1-1* and Col-0 (Figure 6, Supplementary Table 18).

**Figure 6.**
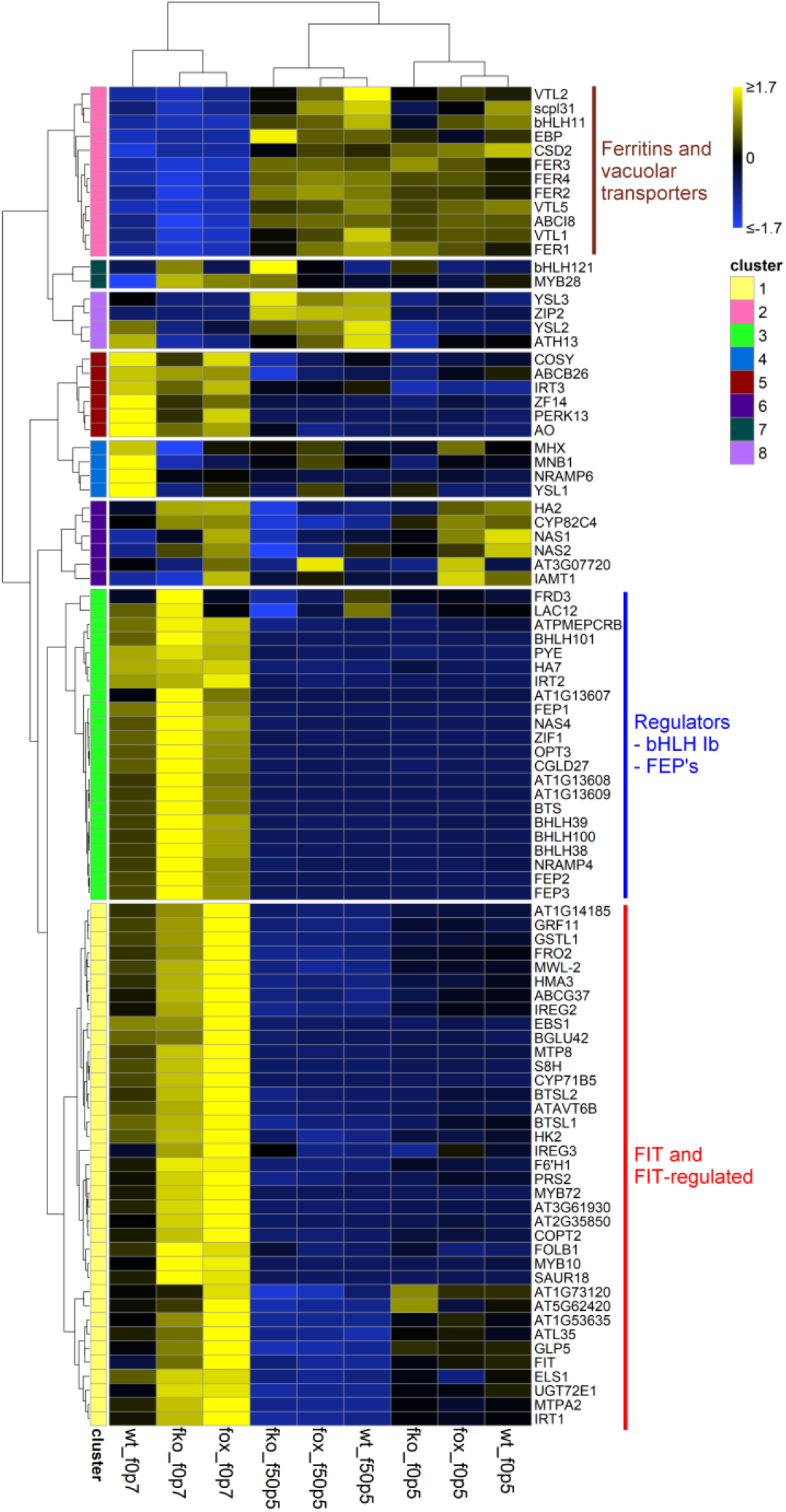
Heatmap of genes related to iron homeostasis as contained in the list “extended 2” from Mai et al., (2023)^22^. These genes were regulated significantly at least once in the pairwise comparisons in our RNA-Seq experiment. Genes from the “extended 2” list that are not included in this heatmap were not found regulated in any of our pairwise comparisons.

In contrast, another cluster was dominated by regulators of iron homeostasis, including the bHLH Ib transcription factors bHLH038, bHLH039, bHLH100, and bHLH101, as well as *FEP1*, *FEP2*, *FEP3*, *BTS*, *PYE*, *NAS2*, *NRAMP4*, *OPT3*, and *FRD3*^23^. These genes were expressed at the highest levels in *fdk1-1* (Figure 6, Supplementary Table 18). Additional clusters showed highest expression in Col-0 under -Fe pH 7.5 and included genes such as *COSY*, *YSL1*, *NRAMP6*, and *IRT3*.

Thus, *FIT*-associated genes and upstream iron-homeostasis regulators displayed contrasting expression patterns across genotypes under alkaline iron deficiency.

## Discussion

### *FDK1* contributes to adaptation to alkaline iron deficiency

Although *FDK1* is strongly induced by iron deficiency, its function has remained unknown. The longer-root phenotype of *fdk1-1* under both iron regimes, and its restoration in *FDK1-Ox*, indicate that *FDK1* contributes to root growth regulation. By contrast, loss of *FDK1* had only modest or inconsistent effects on ferric reductase activity and pigment accumulation, and IAN and camalexin levels were unchanged. *FDK1* therefore does not appear to be an essential component of the canonical Strategy I uptake machinery or a major regulator of the tested indolic metabolites.

FDK1-GFP localized to both the cytoplasm and nucleus in *N. benthamiana* and *A. thaliana* roots, without detectable iron-dependent redistribution. This localization is compatible with metabolic or regulatory functions, although it does not establish either. Despite its evolutionary relationship to NSPs^19^, the absence of genotype-dependent changes in IAN and camalexin argues against a major role in the metabolism of these indolic compounds^15,24,25^. The presence of *IDI* homologs throughout the plant kingdom^14^ suggests that its ancestral functions include regulation of iron-responsive specialized metabolism while the NSPs have diverged as iron-dependent enzymes in specialized metabolism.

### FDK1 promotes iron-mobilizing coumarin responses under alkaline iron deficiency

The clearest physiological effect of *FDK1* was observed in coumarin exudation. Scopoletin levels in culture medium increased in all genotypes under −Fe pH 7.5, whereas fraxetin and esculetin showed strong genotype dependence. Their induction under alkaline iron deficiency was attenuated in *fdk1-1* and strongly enhanced in *FDK1-Ox*. Thus, *FDK1* is not required for the general activation of coumarin secretion but contributes specifically to the accumulation of coumarins in the medium or their release for iron mobilization under alkaline conditions. This distinction is physiologically relevant because coumarins support iron acquisition when reductive Strategy I uptake becomes inefficient^5,26^. Fraxetin is particularly effective at mobilizing poorly soluble Fe³⁺ and can form iron complexes that contribute to plant nutrition^5,6,12^. The reduced fraxetin response in *fdk1-1* therefore connects *FDK1* to a functionally important branch of the alkaline iron-deficiency response, while its strong enhancement in *FDK1-Ox* indicates that elevated FDK1 abundance can promote this pathway.

Because scopoletin, a precursor of fraxetin and esculetin^27^, was still induced in *fdk1-1*, FDK1 is unlikely to control the entire coumarin pathway. It may instead affect later biosynthetic steps, pathway flux, conversion of scopoletin-derived intermediates, or export. This interpretation is consistent with the selective metabolite phenotype and argues against a general defect in phenylpropanoid metabolism. An additional possibility is that *FDK1* influences the selective release or hydrolysis of stored coumarin glucosides, thereby affecting the availability of specific aglycones such as fraxetin and esculetin. Such a mechanism could likewise explain why scopoletin remains inducible while downstream coumarins are selectively affected.

### Transcriptomic evidence links *FDK1* to coumarin metabolism and iron homeostasis

The RNA-seq data independently support the results of metabolite profiling. Genes induced under alkaline iron deficiency were GO term-enriched for coumarin metabolism, iron starvation, pH responses, and other secondary metabolic processes. Across several complementary analyses, “coumarin metabolic process” was repeatedly the most strongly enriched term in gene sets associated with high *FDK1* transcript abundance. It characterized cluster 6, which showed highest expression in *FDK1-Ox* under −Fe pH 7.5, as well as manually filtered sets containing genes highest in *FDK1-Ox* or lowest in *fdk1-1*. This recurrence indicates a robust association between *FDK1* and transcriptional activation of coumarin-related pathways.

Expression patterns of individual pathway genes were consistent with this interpretation. Multiple genes encoding enzymes or transporters involved in coumarin biosynthesis and transport, including members of the *PAL* and *4CL* families, *C3H*, *F6′H1*, *S8H*, *COMT1*, *CCoAOMT*s, *COSY*, *BGLU42*, and *PDR9*, differed between genotypes under −Fe pH 7.5. Because these changes affected several steps, the altered coumarin profiles are unlikely to result from regulation of a single reaction. However, the genes did not all follow an identical pattern, indicating that *FDK1* probably influences the pathway through a broader regulatory or metabolic mechanism rather than by uniformly controlling every component.

*FDK1*-associated gene sets were also enriched for GO terms metal transport and homeostasis, including iron, zinc, and transition-metal transport. *FDK1* may therefore be connected not only to rhizosphere iron mobilization but also to downstream uptake, transport, or redistribution. Nevertheless, mineral concentrations were determined mainly by treatment, with comparatively small genotype effects. The transcriptional differences may thus reflect altered regulation or compensation that does not necessarily translate into large changes in bulk mineral content under the tested conditions.

Clusters preferentially expressed in *fdk1-1* contained several genes related to mitochondrial protein import and metabolism, including *TOM40-2*, *mtOM64*, *TIM17-1*, *TIM23-3*, *TIM44-1*, *MGE1*, *MPC2L*/*NRGA1*, *CoAc2*, and *MTM1* (Supplementary Table 11)^28–34^. This pattern may indicate compensatory mitochondrial remodeling or redox adjustment rather than a specific defect in mitochondrial iron transport. Because mitochondrial activity and organellar metabolite or metal contents were not examined, a direct role of FDK1 in mitochondrial function cannot yet be inferred.

### *FDK1* is associated with *FIT*-dependent iron-deficiency responses

A prominent transcriptomic feature was the contrasting behavior of *FIT*-associated and other iron-homeostasis genes. Under −Fe pH 7.5, *FDK1-Ox* showed high expression of *FIT* and many genes linked to *FIT*-dependent iron acquisition and coumarin metabolism, including *IRT1*, *F6′H1*, *S8H*, *PDR9*, *BGLU42*, *MYB10*, and *MYB72*. This expression pattern parallels the enhanced coumarin exudation phenotype of *FDK1-Ox* and suggests a relationship between *FDK1* abundance and *FIT*-centered responses.

By contrast, bHLH Ib transcription factors, regulators and effectors including *BTS*, *PYE*, *FEP1-3*, *NAS2*, *NRAMP4*, *OPT3*, and *FRD3* were most highly expressed in *fdk1-1*. *FDK1* therefore does not appear to amplify the iron-deficiency response globally. Instead, it is associated with a specific balance between *FIT*-dependent responses and other regulatory branches, possibly reflecting altered iron sensing or compensatory signaling in the knockout.

The direction of causality remains unresolved. *FDK1* is induced by iron deficiency and regulated by *FIT*^3,13^, while *FIT* and many *FIT*-responsive genes are elevated in *FDK1-Ox*. *FDK1* could therefore act downstream of *FIT*, feedback on FIT activity, or participate in a reciprocal regulatory circuit. One possible model is that FDK1 promotes coumarin production or secretion, changing local iron mobilization and thereby influencing signals that sustain iron-acquisition responses. Alternatively, FDK1 may act more directly on regulatory processes controlling coumarin-pathway and *FIT*-responsive genes. Its cytoplasmic and nuclear localization is compatible with both possibilities but does not distinguish them from one another.

The transcriptional and metabolic changes did not produce an obvious shoot phenotype under all growth conditions. Phenospex-based phenotyping did not show significant differences between Col-0, *fdk1-1*, and *FDK1-Ox* on control or alkaline calcareous soil. *FDK1*-dependent effects may therefore be subtle, tissue-specific, conditional, or buffered at the whole-plant level.

### Additional metabolic pathways may contribute under alkaline conditions

Although coumarin metabolism showed the strongest association with *FDK1*, clusters preferentially expressed in Col-0 under −Fe pH 7.5 were enriched for flavone- and flavonol-related processes. Flavonoids can chelate Fe²⁺ and Fe³⁺, participate in redox reactions, and contribute to nutrient-deficiency and rhizosphere responses^35–39^. Their enrichment may therefore reflect additional metabolic adaptations to alkaline iron deficiency.

However, flavonoid concentrations were not measured, and the expression data alone do not demonstrate effects on iron acquisition. These pathways may operate alongside coumarins, but their physiological relevance and relationship to FDK1 remain to be established. The results nevertheless support the broader view that adaptation to iron limitation involves several classes of specialized metabolites rather than a single pathway^10,11^.

### How does *FDK1* link iron mobilization and iron signaling?

Our findings place *FDK1* at the interface between coumarin-mediated iron mobilization and FIT-associated transcriptional responses. Loss of *FDK1* attenuated fraxetin and esculetin induction, whereas overexpression led to enhanced levels and was associated with increased expression of coumarin-pathway and FIT-responsive genes. FDK1 may influence biosynthesis enzymes or transporters, regulatory proteins, or both. Post-translational regulation of phenylpropanoid metabolism, including control of PAL stability by Kelch repeat F-box proteins^40^, represents one possible mechanism of interaction. However, for such a kind of interaction to result specifically in the increase of fraxetin or esculetin production, it is expected to happen with enzymes involved in later steps of the pathway.

Future work should determine whether *FDK1* acts directly within the FIT network or through metabolic feedback. Genetic interaction studies, protein-interaction analyses, and measurements of pathway intermediates, enzyme activities, and iron distribution will be required to distinguish these possibilities. Investigating the mitochondrial and flavonoid-related signatures may also clarify whether they represent secondary compensation or additional FDK1-dependent functions.

In conclusion, FDK1 is a component of the alkaline iron-deficiency response that promotes the induction of specific iron-mobilizing coumarins and is associated with coordinated changes in coumarin metabolism and FIT-dependent gene expression. These findings provide a functional explanation for the strong induction of FDK1 during iron deficiency and establish a framework for determining how metabolic iron mobilization is integrated with transcriptional iron-homeostasis networks.

## Materials and Methods

### Plant materials

We used *Arabidopsis thaliana* wildtype Col-0 (N1092) plants, the At3g07720 knockout mutant *fdk1-1* (SALK_020735, N520735, Col-0 background), the At3g07720 overexpression line *FDK1-Ox* (*p35s::HA_3_-FDK1* in *fdk1-1* background), and the *pFDK1::HA_3_-FDK1* construct in Col-0 background.

### Growth conditions

The seeds were sterilized under constant agitation for 8 min in an aqueous solution of 6 % sodium hypochlorite and 0.1 % Triton X-100. The seeds were subsequently washed 5 times with ddH2O, and then dispersed in 0.1 % plant agar in ddH_2_O. The seeds were stored at 4 °C for 48 h before being sown.

For growth on plates, sterilized seeds were directly sown on ½ Hoagland agar plates with either 50 µM (+Fe) or 0 µM iron (-Fe), and with either pH 5.5 and 5 mM MES, or pH 7.5 and 5 mM HEPES, respectively. Plants were grown for the given number of days before roots were harvested.

For small-scale hydroponic growth, seedlings were pre-grown on ½-strength Hoagland agar plates (+Fe). After two weeks the plants were transferred to a 6-well-plate with each well containing 5 ml liquid ½-strength Hoagland medium without sucrose. Each well was filled with ten seedlings. The medium was with (+Fe) or without iron (-Fe) and buffered to either pH 5.5 with 5 mM MES or pH 7.5 with 5 mM HEPES.

For large scale hydroponic growth, plants were pre-grown on 0.3 % agar in 0.5 ml tubes with a hole in the bottom. After 1 week, when the roots had grown through the agar and out of the hole, the seedlings were transferred to boxes with 400 ml liquid ½-strength Hoagland medium at pH 5.8. 6 seedlings were placed per 400 ml of medium. After 4 weeks in the hydroponic medium and a total of 5 weeks, the medium was replaced by either ½-strength Hoagland medium with 50 µM or 0 µM Fe either at pH 5.5 or pH 7.5. After 1 week and a total of 6 weeks, the roots were harvested.

Plants were grown in growth chambers (CU22L or CU36L3, CFL Plant Climatics) under long day conditions with a 16 h day and 8 h night cycle, with 21 °C, 50 % room humidity and 140 μmol·m^−2^·s^−1^ light during the day, and with 19 °C, 50 % room humidity and 0 μmol·m^−2^·s^−1^ light during the night on tall shelves. Growth on agar plates was conducted equally but with 120 μmol·m^−2^·s^−1^ light on shallow shelves.

For growth in alkaline calcareous soil, seeds were sterilized and grown on +Fe plates for eight days as described before. On the eighth day, they were transferred to either potting soil (control, pH 6.0) or potting soil supplemented with 8 g/l calcium carbonate and 4 g/l sodium bicarbonate (ACS, pH 7.7). ACS plants were watered with sodium bicarbonate solution, control plants with dH2O. Plants were phenotyped using a PlantEye (Phenospex, Heerlen, The Netherlands) and photographed with a camera (α6000, Sony, Minato, Japan) at 28 days after sowing. Blue foil was used to simplify background removal in the PlantEye Software (Phena version 2.0, HortControl version 3.85).

### Generation of *FDK1* overexpression and *pFDK1::FDK1-GFP-HIS_6_* plants

The *FDK1* (At3g07720) CDS was cloned into the pAlligator2 vector^41^ using the Gateway™ cloning technique (Invitrogen, ThermoFisher Scientific), to generate the *p35s::HA_3_-FDK1* fusion construct. The vector was then transferred to *Rhizobium radiobacter* GV3101/pMP90. *R. radiobacter* containing the pAlligator2 plasmid with the fusion construct was used to stably transform *fdk1-1* plants. T1 seeds were selected by GFP fluorescence of the seeds, and positive seeds were propagated. T3 seeds were obtained from T2 plants selected with spectinomycin in the growth media. The presence and location of the T-DNA was verified with TAIL-PCR ^42^ using the T-DNA-specific primers S1 (TGGTCCGATTGAGACTTTTC), S2 (CGACAGTGGTCCCAAAGATGG) and S3 (TAGCTTGCATGCCTGCAGGTC), and the arbitrarily degenerate (AD) primers AD1 (NGTCGASWGANAWGAA), AD2 (TGWGNAGSANCASAGA), AD3 (AGWGNAGWANCAWAGG), AD4 (STTGNTASTNCTNTGC), AD5 (NTCGASTWTSGWGTT), and AD6 (WGTGNAGWANCANAGA). TAIL-PCR #3 products were sequenced.

The *pFDK1::FDK1-GFP-HIS_6_* construct was generated by inserting the *FDK1* genetic sequence including the 1000 nt upstream sequence into the pMDC107 vector^43^ using the Gateway™ cloning technique. The *vector was cloned into Rhizobium radiobacter* C58C1(pTiB6S3ΔT)^H^. *R. radiobacter* containing the pMDC107 plasmid with the fusion construct was used to stably transform Col-0 plants. T1 to T3 plants were selected by PCR and by the GFP fluorescence in the roots under -Fe.

### Measurement of phenolic compounds

The liquid medium was collected after the given amount of time. For semiquantitative determination of phenolic compounds, the fluorescence was measured at 365 nm excitation and 460 nm emission using the Tecan Infinite 200 PRO microplate reader. The concentration of specific coumarins such as scopoletin, fraxetin and esculetin in growth medium was determined by HPLC-ESI-MS based on the procedure described in^9,44^. Samples were injected into an Agilent 1200 Series HPLC equipped with a Chromolith (Merck) RP-18e column (100 x 4.6 mm) using a gradient of water/0.1 % (v/v) formic acid (A) and acetonitrile/0.1 % (v/v) formic acid (B) at a flow rate 1 ml/min: 0-4.9 min (0 - 90% B), 4.9-5.0 min (90 – 100% B), 5.0-6.0 min (100% B). Detection was achieved with a Q-TRAP 3200 mass spectrometer (Sciex), operated in negative mode with the following instrument parameters: curtain gas, 40 psi; collision gas setting, high; ion spray potential, -4500 volts; ESI temperature, 450°C; gas1, 75 psi; gas2, 50 psi; entrance potential, 10 volts; exit potential, -1 volts. Coumarins were quantified based on peak areas from multiple reaction monitoring (MRM) transitions (Supplementary Table 19) and external calibration with standards (Sigma-Aldrich). The results were normalized to the root fresh weights. Each biological replicate was conducted with 10 plants, with three independent replicates analyzed.

### Pigment measurements

Pigments were extracted from fresh leaves in acetone. Pigment concentrations were determined spectrophotometrically following Lichtenthaler (1987)^45^. Absorbance of acetone extracts was recorded at 662, 645, and 470 nm using the Tecan Infinite 200 PRO microplate UV–Vis spectrophotometer. Chlorophyll a, chlorophyll b, and total carotenoid concentrations were calculated according to the equations provided for acetone extracts^45^. The results were normalized to the respective fresh weights. For each biological replicate, pigments were extracted from 100 mg fresh material, with three independent replicates analyzed.

### Ferric reductase activity measurements

Root ferric chelate reductase activity was measured following Le et al. (2016)^46^. Seedlings were rinsed with 0.1 M Ca(NO₃)₂, incubated for 1 h in the dark at room temperature in assay solution (300 µM Ferrozine, 100 µM FeNaEDTA), and Fe²⁺ production was quantified photometrically at 562 nm. Reductase activity was expressed per gram of root fresh weight. For each biological replicate, five plants were used, with three independent replicates analyzed.

### IAN and camalexin contents

Indole-3 acetonitrile (IAN) and camalexin were analyzed by LC-MS as described^47,48^. For each biological replicate minerals were extracted from ca. 50 mg of pooled fresh material, with 4-6 independent replicates analyzed. Separation was done on a C18 XSelect™ HSST3 column, 2.5 µm, 3 mm X 150 mm (Waters, Ireland) on a Dionex 3000 HPLC system (Thermo Scientific, Germany). Mobile phase A of the binary gradient system was 99 % H_2_O + 0.1 % formic acid (FA) and mobile phase B was 99 % methanol + 0.1 % FA. The gradient started at 2 % mobile phase B, and was increased to 100 % B from 1-18 min and then held at 100 % for 4 min. The gradient was returned to 2 % mobile phase B within 1 min and the system was equilibrated at 2 % B for 7 min. The flow rate was set to 0.35 ml/min.

IAN and camalexin were analyzed with high resolution mass spectrometry on a maXis 4G Q-TOF instrument (Bruker Daltonics, Germany) equipped with an electrospray ionization source. The operating conditions were as described^48^. IAN was analyzed in negative ionization mode, whereas camalexin was analyzed in positive ionization mode. Identification was done by comparison with standards obtained for indole-3 acetonitrile (OlChemim Ltd., Czech Republic) and camalexin (Sigma-Aldrich, Germany). Quantification was based on calibration curves in relation to the deuterated internal standard ^2^H_5_-indole-3-acetic acid (OlChemim Ltd., Czech Republic), which was used in both negative and positive ionization mode. The internal standard was added to all samples at a concentration of 150 nM during the extraction process.

### Mineral contents

Samples were dried at 65 °C for at least 3 d, ground to a fine powder, and transferred to metal-free 5 mL screw-cap tubes. 1 ml of 67 % nitric acid was added, and samples were digested by boiling until the solution became clear. The solution was centrifuged (5 min, 21,000 × g) and the supernatant was transferred to another tube for analysis. Elemental concentrations were determined at the Biocenter-MS at the University of Cologne by inductively coupled plasma mass spectrometry (ICP-MS; Agilent 7700) following the manufacturer’s instructions, and normalized to sample dry weight^49^. For each biological replicate minerals were extracted from ca. 20 mg of dried material, with five independent replicates analyzed.

### Transcriptome analysis

Briefly, plants were grown on ½-stength Hoagland agar with either 50 µM at pH 5.5, 0 µM Fe at pH 5.5, or 0 µM Fe at pH 7.5. After 9 days, the roots were harvested, and total RNA was extracted using the RNeasy Plant Mini Kit (Qiagen). The libraries were sequenced as 101-bp single-end reads using Illumina HiSeq at the Biologisch-Medizinisches Forschungszentrum (Heinrich Heine University, Düsseldorf, Germany). The reads were then analyzed using the PlugNSeq pipeline^50^.

Functional enrichment analysis of the differentially expressed genes was performed using the clusterProfiler R package^51^. Gene Ontology (GO) terms for *A. thaliana* were sourced from the org.At.tair.db genome-wide annotation database^52^. Querying and mapping of gene identifiers were facilitated using the AnnotationDbi data interface package^53^. All statistical analyses and visualizations were conducted using R version 4.6. Each biological replicate consisted of 100 mg of pooled fresh material, with three independent replicates analyzed.

## Acknowledgements

The authors thank Sabine Ambrosius of the Biocenter-MS at the University of Cologne for her contribution in ICP-MS (minerals) and LC-MS (IAN, camalexin) measurements. The project received funds from the Strategic Research Fund (SFF) of the Heinrich Heine University Dusseldorf, granted to HJM (SFF, F 2017/1259-4). The work was funded by the Deutsche Forschungsgemeinschaft (DFG, German Research Foundation) grant - Project ID 456082119 - TRR 341/1 project A08 to P.B. This work was funded by the Deutsche Forschungsgemeinschaft (DFG, German Research Foundation) under Germanýs Excellence Strategy – EXC-2048/1 – project ID 390686111.

## Author Contributions

Conceptualization (PB, HM), Methodology (HM, PB), Investigation (CvdM, JCL, LS, ES, MCK, MS, VW, HM), Data Analysis (HM, MS, VW), Writing - Original Draft (HM), Review & Editing (all authors), Supervision (HJM, PB, UW), Funding Acquisition (HJM, PB).

## Data availability statement

The transcriptome data are available through GEO accession GSE339092.

## Additional information

The authors declare that there are no competing interests.

## Supplementary Figures

**Supplementary Figure 1.**
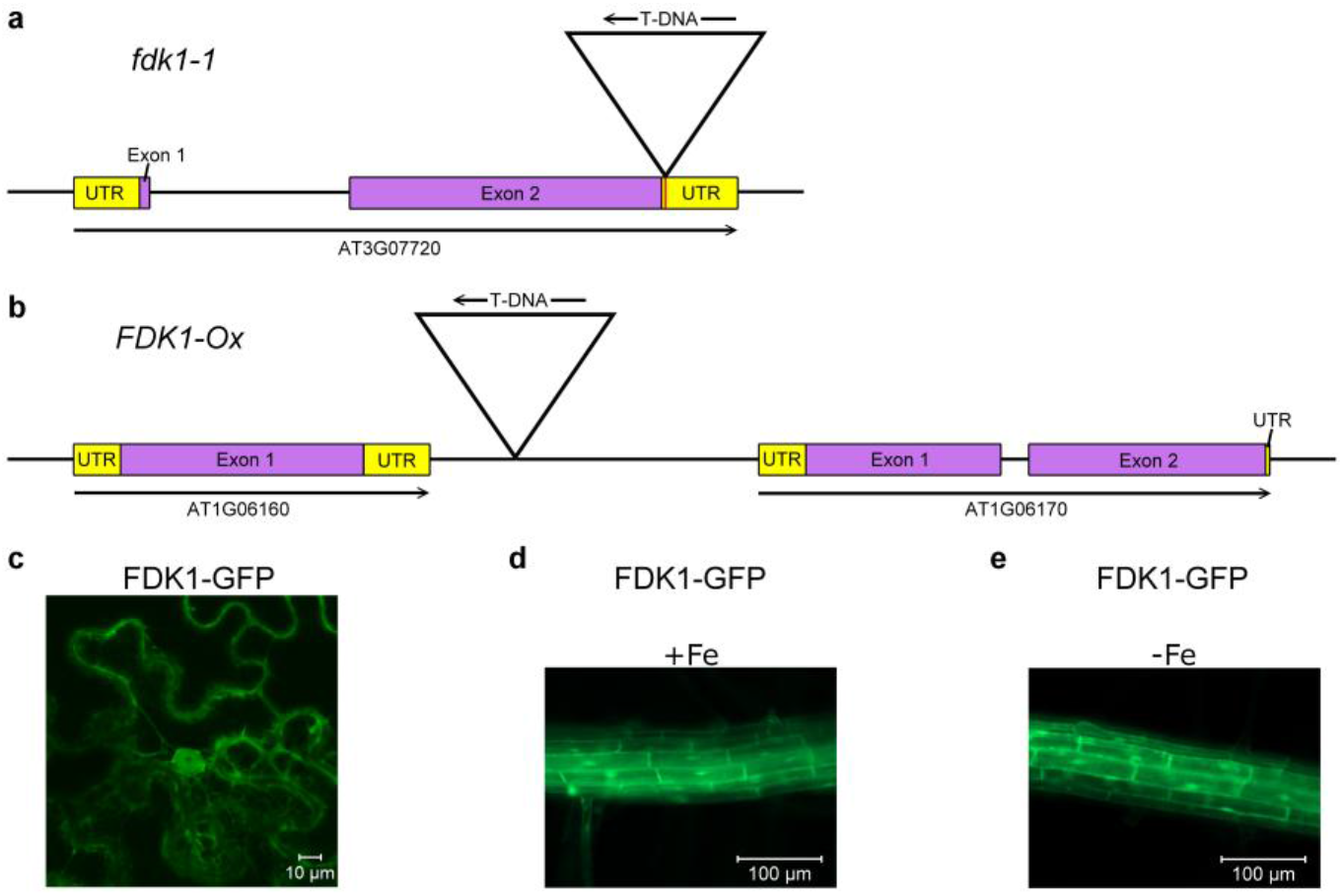
Insertion sites of the FDK1 mutants used in this analysis, and localization of FDK1-GFP in tobacco leaves and Arabidopsis roots. a: insertion site of the T-DNA in the 3’-Utr of AT3G07720 in *fdk1-1* knockout mutants, b: insertion site of the T-DNA in the intergenic region between AT1G06160 and AT1G06170 in *FDK1-Ox* plants, c: localization of FDK1-GFP in the cytosol and the nucleus in tobacco leaves, d: localization of FDK1-GFP in the cytosol and nucleus in Arabidopsis roots under +Fe, e: localization of FDK1-GFP in the cytosol and the nucleus in Arabidopsis roots under -Fe.

**Supplementary Figure 2.**
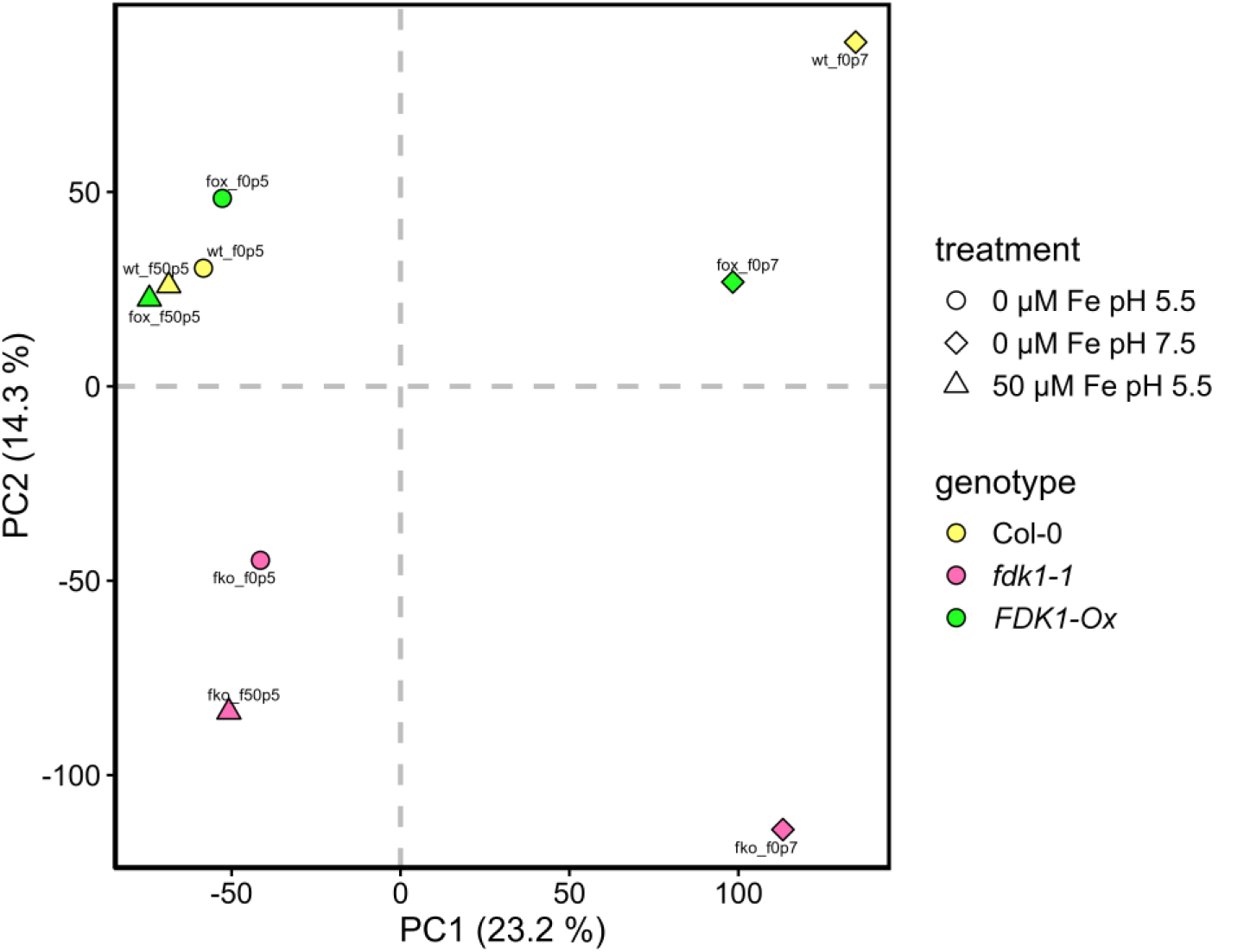
PCA of sample group means of our RNA-seq experiment. Genotypes and treatments have been abbreviated to keep the in the graph labels short. Genotypes are: wt = wildtype, fko = *fdk1-1* knockout mutant, fox = *FDK1-Ox* overexpression mutant. Treatments are: f50p5 = 50 µM Fe pH 5.5, f0p5 = 0 µM Fe pH 5.5, f0p7 = 0 µM Fe pH 7.5.

**Supplementary Figure 3.**
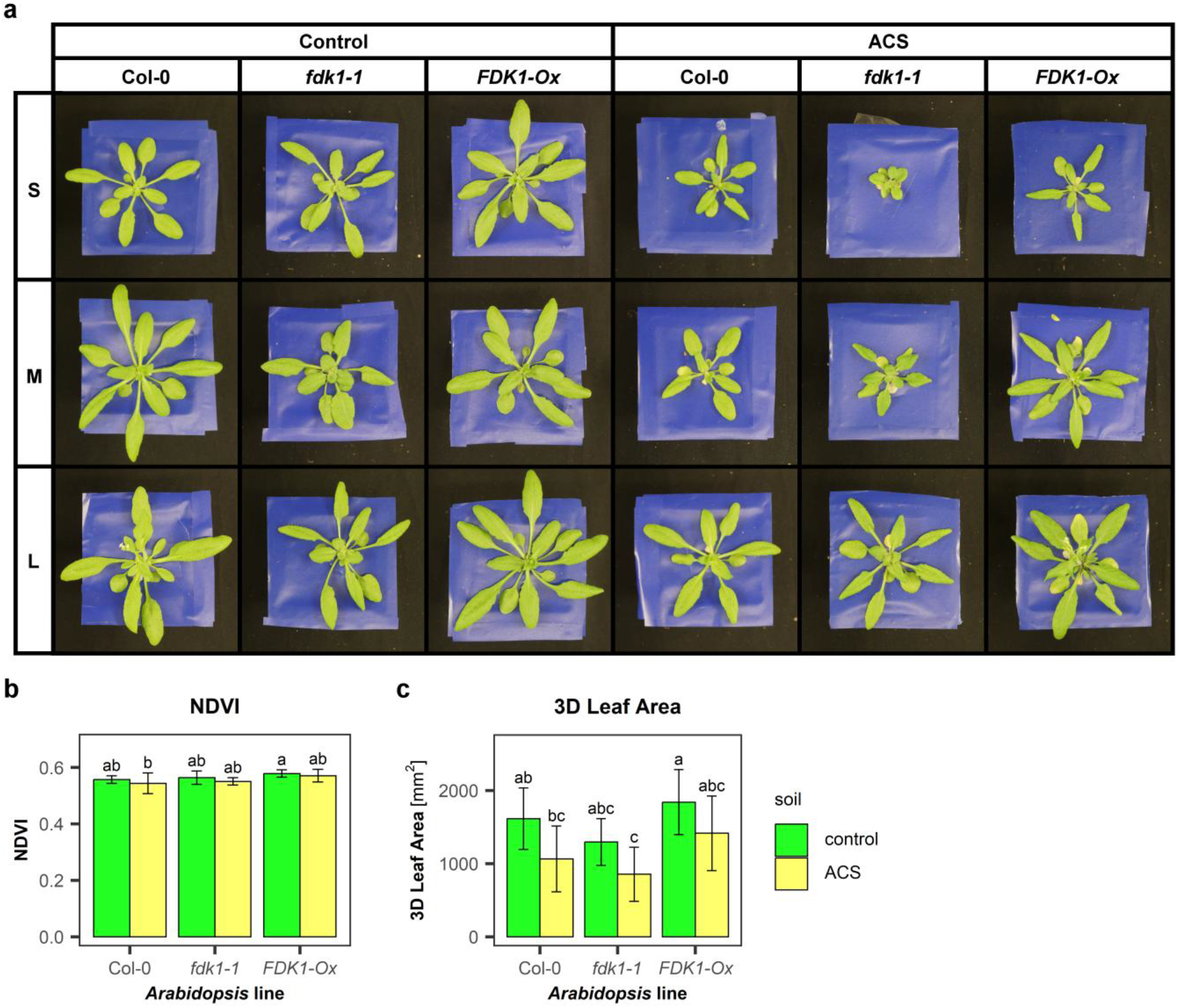
Selected phenotypes of wildtype Col-0, *fdk1-1* and *FDK1-Ox* plants grown on control soil or on alkaline calcareous soil (ACS). a: photos of 28-day-old plants grown on control soil or ACS, respectively. S, M and L denote a small, medium and large specimen of the sample to show the sample variance, respectively. b: NDVI measured by Phenospex. c: 3D Leaf Area measured by Phenospex.

## Notes

### Competing Interest Statement

The authors have declared no competing interest.

